# Visualizing the Coordination of APE1 and DNA Polymerase β During Base Excision Repair

**DOI:** 10.1101/2022.05.02.490312

**Authors:** Max S. Fairlamb, Todd M. Washington, Bret D. Freudenthal

## Abstract

Base Excision Repair (BER) is carried out by a series of DNA repair proteins that function in a step-by-step process to identify, remove, and replace DNA damage. As DNA damage is processed during BER, the DNA transitions through various intermediate states, called BER intermediates, which if left exposed can develop into double-strand DNA breaks and trigger programmed cell death signaling. Previous studies have proposed that in order to minimize exposure of the BER intermediates, each protein may remain bound to its product prior to the next protein binding. Thus, a short-lived complex consisting of the BER intermediate, the incoming enzyme, and the outgoing enzyme may form between each step of the BER pathway. The transfer of BER intermediates between enzymes, known as BER coordination, has yet to be directly visualized and the mechanistic details of the process remain unclear. Here, we utilize single-molecule total internal reflection fluorescence (TIRF) microscopy to investigate the mechanism of BER coordination between apurinic/apyrimidinic endonuclease 1 (APE1) and DNA polymerase β (Pol β). When preformed complexes comprised of APE1 and the incised AP-site product were subsequently bound by Pol β, the Pol β enzyme dissociated shortly after binding in a majority of the observations. In the events where Pol β binding was followed by APE1 dissociation (i.e., DNA hand-off), Pol β had remained bound for a longer period of time to allow disassociation of APE1. Our results indicate that, in the absence of other BER factors, transfer of the BER intermediate from APE1 to Pol β during BER is dependent on the dissociation kinetics of APE1 and the duration that Pol β remains bound near the APE1-5’ nick complex. These findings provide insight into how APE1 and Pol β coordinate the transfer of DNA within the BER pathway.

## INTRODUCTION

It’s estimated that the DNA in each human cell is damaged over 10,000 times every day as a result of exogenous factors, such as exposure to radiation, and endogenous factors, such as the reactive oxygen byproducts of metabolism (1-4). To repair oxidative DNA damage, human cells utilize a multi-step DNA repair pathway called base excision repair (BER) (5). BER is initiated when a damaged nucleobase is bound by a DNA glycosylase, which cleaves the N-glycosidic bond of the damaged nucleotide base, resulting in a baseless sugar entity called an apurinic/apyrimidic (AP) site (Figure 1). This AP site is a substrate for human apurinic/apyrimidinic endonuclease 1 (APE1), which incises the DNA backbone on the 5’ side of the AP site. The resulting product, herein referred to as 5’ nick, consists of a single nucleotide gap and a 5′-deoxyribose phosphate flap (dRP) flap. The 5’-dRP flap is excised by DNA polymerase beta (Pol β), resulting in a 1-nucleotide gap intermediate called the 1-nt gap. Pol β also inserts a new nucleotide across from the orphaned base, resulting in a single stranded nick on the 3’ side of the newly inserted nucleotide, referred to as the 3’ nick. Finally, the last enzymatic step is performed by the X-ray cross-complementing group 1 and DNA ligase III alpha heterodimer (XRCC1:Lig3α), which seals the single stranded nick and completes the BER pathway.

**Figure 1.**
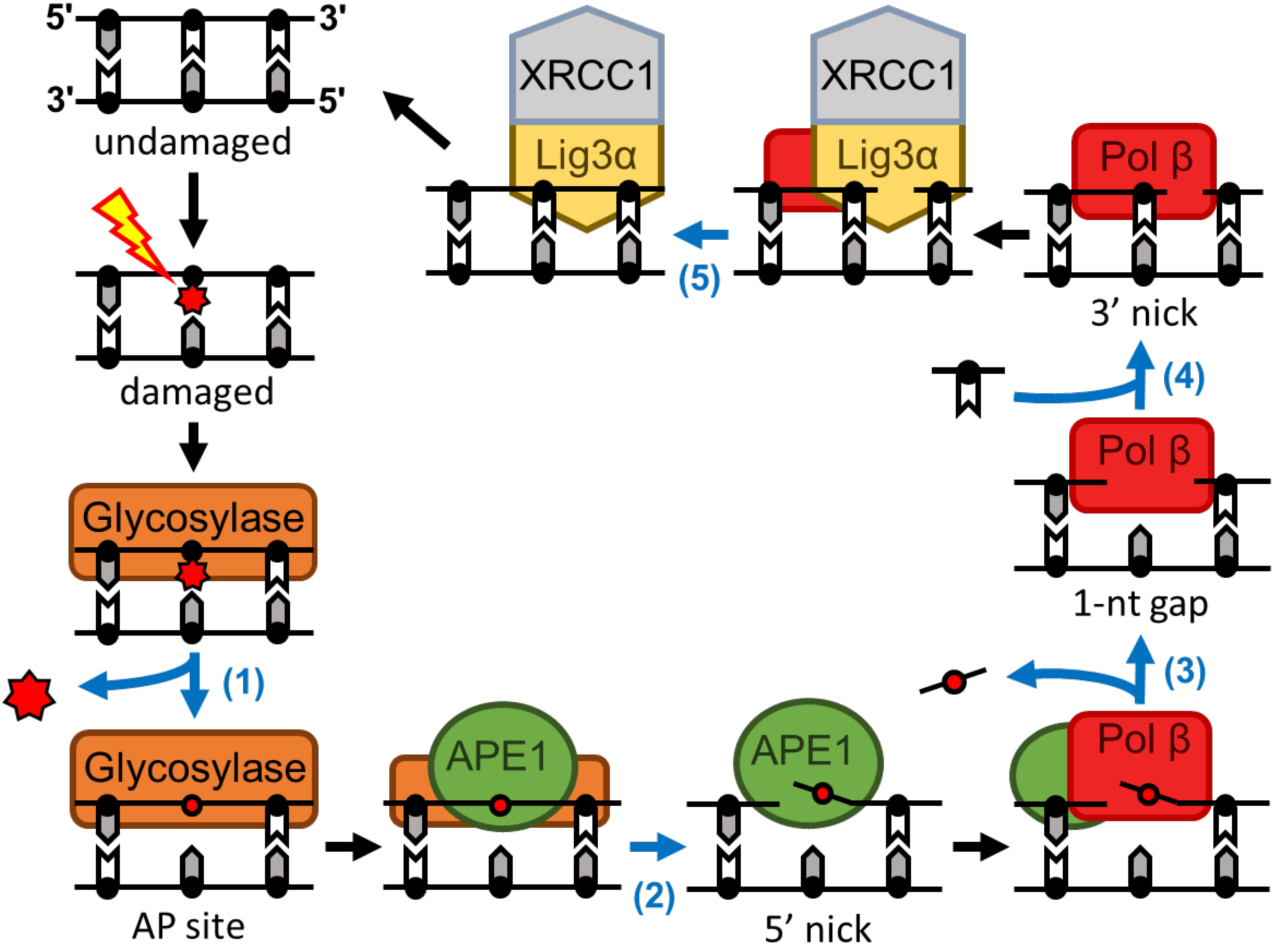
Base excision repair pathway. Base excision repair pathway consists of five catalytic steps (blue arrows). After the formation of DNA base damage, a DNA glycosylase cleaves the N-glycosidic bond of the damaged base (1) generating an AP site. APE1 cleaves the DNA backbone on the 5’ side of the AP site (2), resulting in a 5’ nick substrate, which consist of a single nucleotide gap and a baseless 5′-deoxyribose phosphate (dRP) flap. Pol β excises the dRP flap (3), resulting in a 1-nt gap substrate, and inserts a nucleotide across from the templating base (4), resulting in the 3’ nick substrate. Repair is ultimately completed by the XRCC1:Lig3α heterodimer which ligates the single stranded nick (5). The BER coordination model proposes that a short-lived ternary complex consisting of the DNA intermediate, the incoming enzyme, and the outgoing enzyme forms between each catalytic step of BER. These ternary complexes are depicted between the catalytic steps of the pathway (before 2, before 3, and before 5).

The various intermediate states that the DNA damage transitions through during BER (AP site, 5’ nick, 1-nt gap, 3’ nick) are referred to as BER intermediates. If BER intermediates are left unrepaired, they can degrade into double-stranded DNA breaks, inhibit the progression of replicative DNA polymerases and/or trigger programmed cell death signaling (6-8). To stabilize the BER intermediates during repair, it has been proposed that BER enzymes remain bound to their respective BER intermediate product until the next enzyme in the pathway binds the DNA intermediate (9-11). This transfer of BER intermediates between BER enzymes is referred to as BER coordination, and has been described as analogous to runners passing a baton (12). The importance of this mechanism is highlighted by previous studies which have shown that disruption of BER coordination can result in diminished BER efficiency and genomic instability (13,14).

Consistent with the model of BER coordination, previous kinetic and protein footprinting studies indicate that transfer of the 5’ nick DNA substrate from APE1 to Pol β involves the formation of a three-member complex composed of APE1, Pol β, and the 5’ nick BER intermediate (11,15,16). We refer to this transient formation as the APE1-Pol β-5’ nick ternary complex. The details of how APE1-Pol β-5’ nick ternary complex formation promotes transference of the 5’ nick BER intermediate between APE1 and Pol β remain unclear. Part of the challenge in characterizing this BER coordination mechanism has been the difficulty of resolving the timing of association and dissociation steps that occur during the formation of APE1-Pol β-5’ nick ternary complexes. For example, Prasad et al. demonstrated that an AP site could be incised by APE1, transferred to Pol β, and further processed into downstream BER intermediates within a 10 second time frame. However, these experiments were unable to resolve how long the APE1-Pol β-5’ nick ternary complex remained intact or the order of APE1 and Pol β assembly and disassembly from the BER intermediate. These details could provide critical insight into the mechanism of BER coordination between APE1 and Pol β.

Here, we utilized single-molecule total internal reflection fluorescence (TIRF) microscopy to visualize the sequence and timing of association and disassociation steps that occur during the formation of APE1-Pol β-5’ nick ternary complexes. Our results indicate that APE1 and Pol β both form the most stable interactions with their respective substrates and products relative to other BER intermediates in the pathway. We also find that the association of Pol β to a preformed APE1-5’ nick complex was most frequently followed by rapid Pol β dissociation. In the majority of events where Pol β association was followed by APE1 dissociation, the APE1-Pol β-5’ nick ternary complex was longer-lived. Based on these findings, we propose that in the absence of other BER factors, DNA hand-off events involving APE1 and Pol β reflect the probability that APE1 product release will occur before Pol β dissociates from the complex.

## MATERIAL AND METHODS

### DNA substrates

Oligonucleotides were obtained from IDT. Annealing reactions were performed by mixing the oligonucleotides in a 1:1.2 ratio of biotinylated template to unbiotinylated complement(s). Reaction mixtures were heated to 95 °C for 5 minutes and then allowed to cool slowly to 4 °C. To generate the 25-mer duplex DNA used to capture APE1-DNA and Pol β-DNA binary complex formations, a 5’-biotin end-labeled 25-mer template (5’-[biotin]TGT GTG GAA TAC AGT GAG CGC AAC G-3’) was annealed with the respective oligo(s) described below to yield duplex substrates that mimicked each of the canonical BER intermediates. The AP site BER intermediate was prepared from the template and a complement sequence that contained a centrally placed tetrahydrofuran (THF) group (5’-CGT TGC GCT CA[THF] TGT ATT CCA CAC A-3’). The 5’ nick BER intermediate was prepared from the template, a downstream 11-mer primer (5’-CGT TGC GCT CA-3’), and an upstream 13-mer primer containing a 5’ deoxyribose phosphate group (dRP) and terminal phosphate (phos) (5’-[phos][dRP]TGT ATT CCA CAC A-3’). The 1-nt gap BER intermediate was generated by the same approach using an upstream 13-mer primer which lacked the 5’-dRP (5’-[phos]TGT ATT CCA CAC A-3’). The 3’ nick BER intermediate was generated using the same upstream 13-mer primer and a downstream 12-mer primer (5’-CGT TGC GCT CAC-3’). A 3’-Cy5-labeled version of the 25-mer template was used to collect binary events between Pol β and AP site DNA. The 40-mer immobilized DNA substrate used for capturing APE1-Pol β-5’ nick ternary complex formations was prepared from a 5’-biotinylated and 3’-Cy5 labeled template (5’-[biotin]ATG CAT GTT GTG TGG AAT ACA GTG AGC GCA ACG CAA TCA C[Cy5]-3’), a 21-mer downstream primer (5’-[phos][dRP]T GTA TTC CAC ACA ACA TGC AT-3’), and an 18-mer upstream primer (5’-GTG ATT GCG TTG CGC TCA-3’).

### Protein purification and fluorescent labeling

Human wildtype APE1 and wildtype Pol β were expressed and purified as previously described (17,18). Briefly, protein was overexpressed in BL21(DE3)plysS *E. coli* cells (Invitrogen) bearing a pET28a plasmid (Genscript) that encoded either APE1 or Pol β. Cells were grown in 2xYT at 37°C, expression was induced with IPTG, and harvested cells were lysed via sonication. Protein was purified from the cell lysate via heparin, cation exchange, and gel filtration resins using an FPLC (ATKA-Pure). Purity and concentration of the resulting protein solution was confirmed by SDS-PAGE analysis and by absorbance at 280 nm.

APE1 and Pol β were fluorescently labeled with Cy3 and Cy5 NHS ester dye respectively. APE1 was dialyzed in labeling buffer containing 50 mM HEPES, pH 7.05, and 150 mM sodium chloride and concentrated to 15 mg/mL. While avoiding exposure to light, an Amersham Cy3 mono-reactive NHS ester dye pack (GE healthcare) was resuspended with labeling buffer and mixed with enough purified APE1 to yield a 100 μL solution of 10 mg/mL APE1. The reaction was rotated slowly in darkness for 16 h at 4°C. Excess dye was removed by running the labeling reaction over a cation exchange column using an FPLC (ATKA-Pure). Fluorescently labeling of Pol β followed the same procedure using an Amersham Cy5 mono-reactive NHS ester dye pack (GE healthcare). Labeling efficiency was determined by calculating the molar ratio of dye-to-protein within the solution by measuring absorbance at 280 nm and 530 nm for APE1-Cy3 or 649 nm for Pol β-Cy5. For both Cy3-labeled APE1 and Cy5-labeled Pol β, labeling efficiencies were between approximately 2.1-2.4 labels per protein. Single use aliquots of fluorescently labeled proteins were flash-frozen with liquid nitrogen and stored until use at −80°C for up to 1 month.

### Single molecule imaging and data collection

The single molecule data was collected on a prism-type total internal reflection fluorescence (TIRF) microscope which has been previously described (19). Briefly, excitation lasers (OBIS Coherent) with wavelengths of 532 and 640 nm are merged using a series of dichroic mirrors (Shamrock) and directed to the stage of an inverted microscope (Olympus America, Inc.) using broadband mirrors (Thorlabs). The 532 and 640 nm excitation lasers were operated at 60 mW and 30 mW power respectively. The excitation beams pass through a Pellin-Broca prism (Eksma Optics) which modulates the trajectory of the beams and generates an evanescent field of excitation within the visualized region of the sample chamber. Cy3 and Cy5 emissions were observed using a 60x objective lens (Olympus) and split using an Optosplit-III emission image spitter (Cairn). Recordings were captured using an IXON ULTRA 897 EMCCD camera (Andor) at ten frames per second.

TIRF experiments were performed using quartz sample chambers coated with Biotin-PEG-SVA (Laysan Bio) which were cleaned and prepared as previously described (20,21). The sample chamber was first flushed with 0.2 mg/mL neutravidin (ThermoFisher) in buffer (50 mM HEPES, pH 7.4, 150 nM NaCl) and allowed to sit for 3 min, then unbound neutravidin was flushed out with buffer. All subsequent steps used imaging buffer containing 50 mM HEPES, pH 7.4, 150 mM sodium chloride, 0.1 mg/mL BSA, 0.8% glucose, and 2.5 mM EDTA. For fluorophore longevity and stability, imaging buffer also contained approximately 10 mM Trolox (Cayman), 0.04 mg/mL Catalase (Sigma-Aldrich), and 1 mg/mL Glucose Oxidase (Sigma-Aldrich) as previously described (22). Unless otherwise stated, binary protein-DNA events were collected by first incubating a 350 pM solution of 25-mer DNA in the sample chamber for 3 min, flushing the unbound DNA, and then flowing 0.1 nM of each fluorescently labeled protein into the chamber. Binding events between proteins and immobilized DNA were recorded for 20 min at room temperature and fluorescent trajectories were extracted from the recordings. Experiments designed to capture the formation of APE1-Pol β-5’ nick ternary complexes were performed similarly, except a 15 pM solution of Cy5-labeled 40-mer DNA was incubated in the sample chamber. Also, after starting the recording, the Cy5-labeled DNA was bleached by flushing the chamber with buffer lacking the oxygen scavenging system before adding 1.5 nM of each fluorescently labeled protein.

Fluorescence trajectories, which display the measured intensity of each fluorophore at each immobilized DNA substrate as a function of time, were generated from recordings as previously described (22). Sustained increases in fluorescence intensity which reflected APE1-Cy3 or Pol β-Cy5 binding events to immobilized DNA and satisfied the selection rules as defined in (23), were selected for analysis and idealized with QuB (24) (University of Buffalo). For experiments that utilized Cy5-labeled DNA, only the fluorescence trajectories that exhibited Cy5 signal from the beginning of the recording were analyzed. Events were then organized into an event catalog using the Kinetic Event Resolving Algorithm (KERA) software package, which yields the relative frequency of each type of event observed and the duration between each association and disassociation step within each event (25).

### CRTD analysis

To quantify the dissociation kinetics of a particular complex formation, the lifetimes of all observed complexes (i.e., the interval of time between the association and dissociation steps) were compiled into a cumulative residence time distribution (CRTD) plot (26). The CRTD plot can be interpreted as a type of normalized survival curve which describes the fraction of objects remaining as a function of time when the start of all events is synchronized to time = 0. The probability of observing a protein-DNA complex with a lifetime (*T*) that is greater than or equal to a given time (*t*) decreases exponentially as time *t* is increased. This can be described by

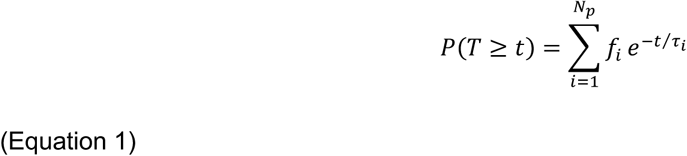

where P(T ≥ t) is the probability that the duration of a randomly selected event will be at least as long as time *t*. The term *N*_*p*_ denotes the number of distinct Poisson processes which contribute to the dissociation kinetics of the overall population, and *f*_*i*_ represents the relative fraction of all observed objects represented by each subpopulation (*i*). The term *τ*_i_ is the average lifetime of the complex, which can be used to derive the dissociation rate constant k_d,i_ (*τ*_i_ = k_d,i_^−1^). To determine the minimum number of Poisson processes that adequately captures all of the features of a given dataset, CRTD plots were fit to single (*N*_*p*_ = 1), double (*N*_*p*_ = 2), and triple (*N*_*p*_ = 3) exponential models and the most likely *N*_*p*_ value was determined via residual analysis and the extra sum-of-squares F test performed using GraphPad Prism version 8.2.1 for Windows (GraphPad Software, La Jolla California, USA) (28). From this analysis, we found that each dataset analyzed was best described by either a double or triple exponential model.

## RESULTS

### APE1 and Pol β binary DNA interactions with BER intermediates

To understand how the state of the BER intermediate affects APE1 binding specificity, we measured the duration that APE1 remained bound to each BER intermediate using single-molecule TIRF microscopy (Figure 2). We refer to binding events which consist of one protein and one DNA molecule as binary events. In each experiment, fluorescently labeled protein was flowed into a sample chamber containing immobilized DNA that mimicked a specific BER intermediate (AP site, 5’ nick, 1-nt gap, 3’ nick, undamaged) and binary events were recorded. Fluorescent trajectories, which depict the fluorescence intensity observed at a single immobilized DNA molecule over as a function of time (Figure 2), were then extracted from the recordings. The duration that each binary complex remained intact (i.e., the complex lifetime) was compiled into a cumulative residence time distribution (CRTD) plot, which represents the percent of observed protein-DNA complexes that remained bound as a function of time (26).

**Figure 2.**
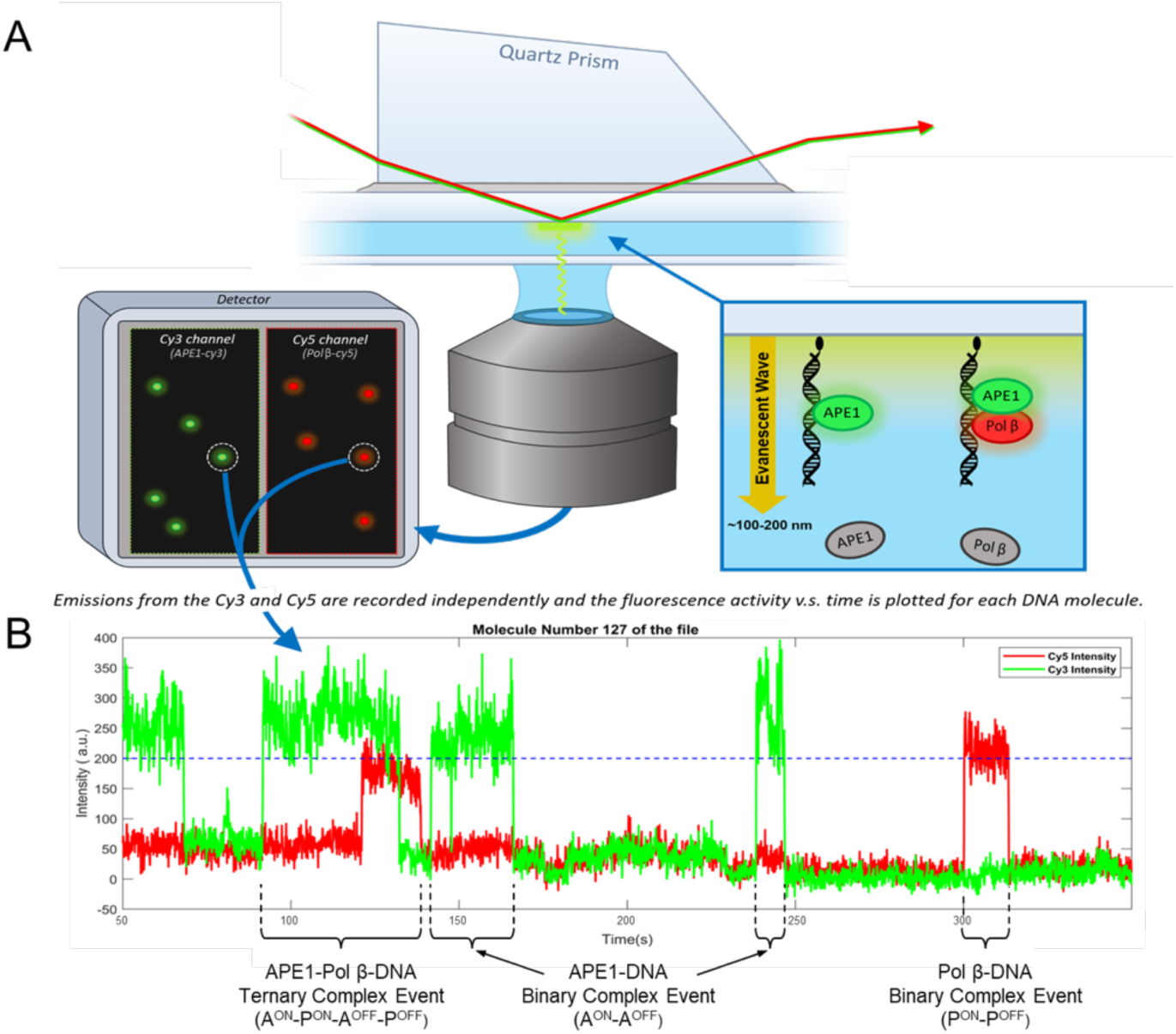
Single molecule TIRF experimental setup. **(A)** 532 nm and 640 nm excitation lasers (red and green arrows) are directed to the imaged region of the sample chamber via a quartz prism. Total internal reflection of the excitation beams produces an evanescent wave (yellow arrow) which excites fluorophores located within ∼100-200 nm of the slide surface. Emission from excited APE1-Cy3 (green oval) and Pol β-Cy5 (red oval) molecules is recorded at 10 frames per second using an EMCCD. **(B)** Fluorescence trajectories depict the fluorescence intensity recorded at an immobilized DNA substrate over time. The association of APE1-Cy3 or Pol β-Cy5 with an immobilized DNA molecule is indicated by an increase in fluorescence intensity in the Cy3 or Cy5 channel above the baseline, respectively, while dissociation is indicated by a decrease in intensity back to baseline. Binary complex events involve association and dissociation of a single protein, either APE1-Cy3 or Pol β-Cy5, from the DNA substrate. Ternary complex events involve the association of both APE1-Cy3 and Pol β-Cy5, resulting in the formation of a three-member complex, before the dissociation of protein.

The CRTD plots describing the dissociation kinetics of binary complexes between APE1 and each of the BER intermediates are presented in Figure 3 A-E. Each CRTD plot was fit to an exponential decay model to yield the mean lifetime of the complex. For each interaction tested, the features of the CRTD plot were best described by a multiexponential decay corresponding to either two or three Poisson processes depending on the BER intermediate tested (Supplemental Table 1). This indicates the existence of multiple independent subpopulations of APE1-DNA binding events within each dataset, which we denote as T_1_, T_2_, and when applicable T_3_. The population of molecules that dissociated according to the short-lived T_1_ process had an average lifetime between 1.2-3.0 sec for each BER intermediate analyzed (Supplemental Table 1). These short T_1_ populations likely reflect interactions that are independent of the intermediate state of the DNA. The timescale of the longer-lasting T_2_ and T_3_ populations were dependent on the BER intermediate being tested, suggesting that these population of events relate to engagement of the DNA lesion. APE1 exhibited the longest average lifetime on the AP site BER intermediate (T_2_ = 41.1 sec, Figure 3A). Meanwhile, APE1 dissociated faster from the 5’ nick BER intermediate (T_2_ = 8.3 sec; T_3_ = 36.5 sec) and the downstream BER intermediates, 1-nt gap (T_2_ = 4.3 sec; T_3_ = 30.9 sec) and 3’ nick (T_2_ = 12.4 sec) (Figure 3B-D). Although the T_2_ population of APE1-undamaged DNA complexes averaged 22.8 sec, the majority of APE1-undamaged DNA complexes followed the shorter-lived T1 process (63.7%) with a mean lifetime of 2.6 sec (Figure 3E). These findings suggest that the binding specificity of APE1 to the DNA lesion could promote APE1 association at the correct stage of BER.

**Figure 3.**
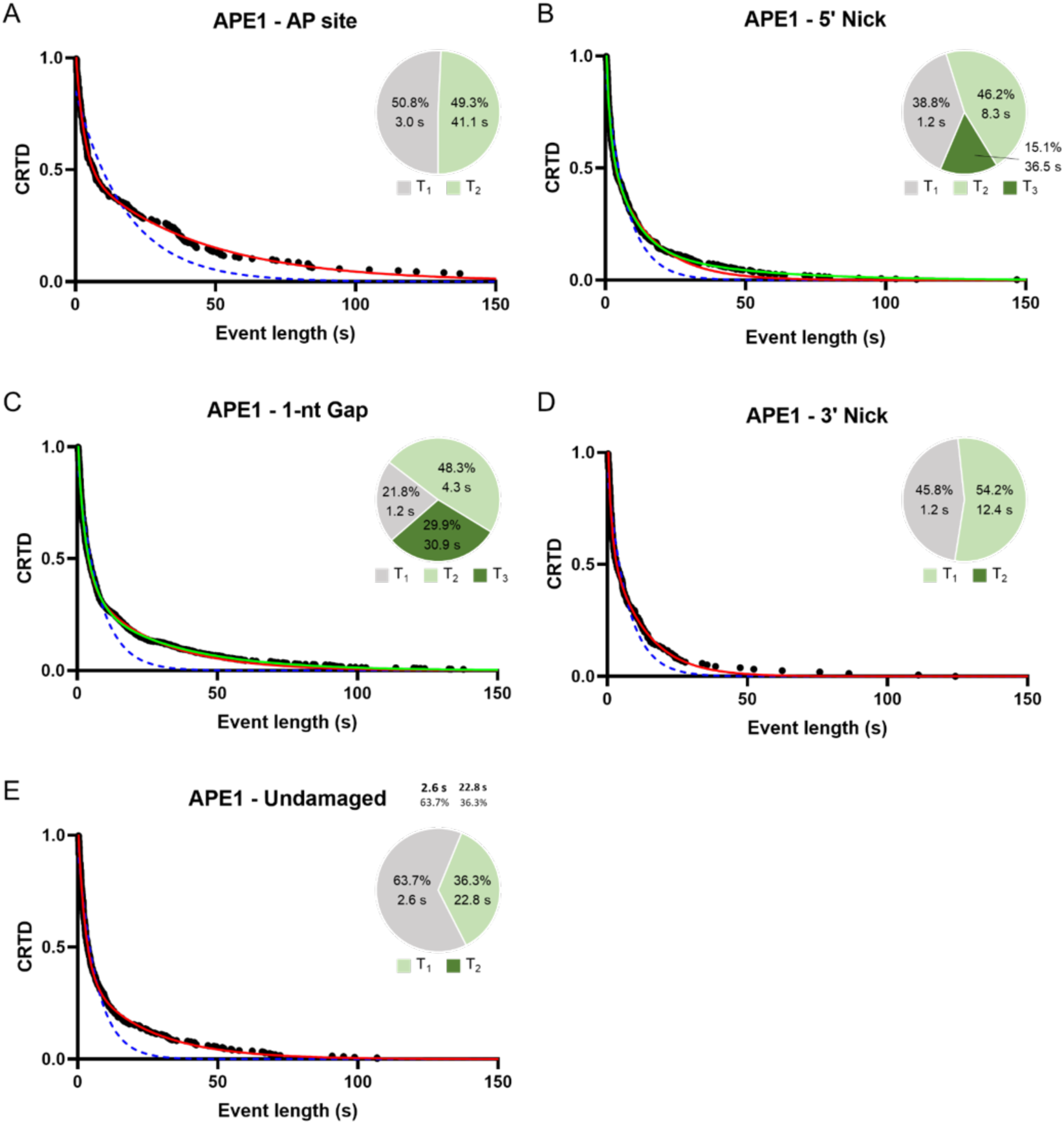
APE1-DNA binary interactions. CRTD vs. time plot of binary complex lifetimes measured from APE1 binding events with DNA substrates containing an AP site **(A)**, 5’ nick **(B)**, 1-nt gap **(C)**, 3’ nick **(D**), and no DNA damage **(E)**. Experimental data is represented by black circles. Single, double, and triple exponential decay fits are depicted as dashed blue, solid red, and solid green lines respectively. Pie charts show the relative fraction of molecules associated with the T_1_, T_2_, and if applicable T_3_ populations. The kinetic parameters and 95% confidence intervals described by the fits are shown in Supplemental Table 1.

Binary Pol β-DNA binding events for each BER intermediate were characterized similar to the APE1-DNA interactions described above. Initial attempts to record binary interactions between Pol β and DNA containing an AP site or undamaged DNA resulted in very few stable complexes (Figure 4A). Consistent with previous findings, this suggests that these intermediates are not preferred substrates of Pol β binding due to the lack of a free 5’ phosphate (16). In contrast, interactions between Pol β and DNA containing 5’ nick, 1-nt gap, and 3’ nick were readily observed (Figure 4B-D). These datasets were best described by either two or three kinetic processes depending on the BER intermediate tested (Supplemental Table 2). Binding interactions between Pol β and the 1-nt gap substrate exhibited a T_2_ mean lifetime of 19.8 sec. Meanwhile, binding interactions between Pol β and the 5’ nick exhibited a T_2_ and T_3_ mean lifetime of 5.2 sec and 28.6 sec and binding interactions between Pol β and the 3’ nick exhibited a T_2_ and T_3_ mean lifetime of 7.4 sec and 33.7 sec. Analyzing the Pol β-AP site complexes indicated they are short-lived, with a mean T_2_ lifetime of approximately 0.9 sec. Altogether, these data indicate that Pol β-DNA binding is highly selective for BER intermediates which contain single stranded nicks (5’ nick, 1-nt gap, and 3’ nick). The short-lived interactions between Pol β and undamaged DNA or DNA containing an AP site suggests a lack of lesion engagement by Pol β, resulting in only fast dissociating events being observed. These results suggest that the specific DNA binding properties of Pol β may help ensure Pol β associates with the DNA at the correct stage of the BER pathway.

**Figure 4.**
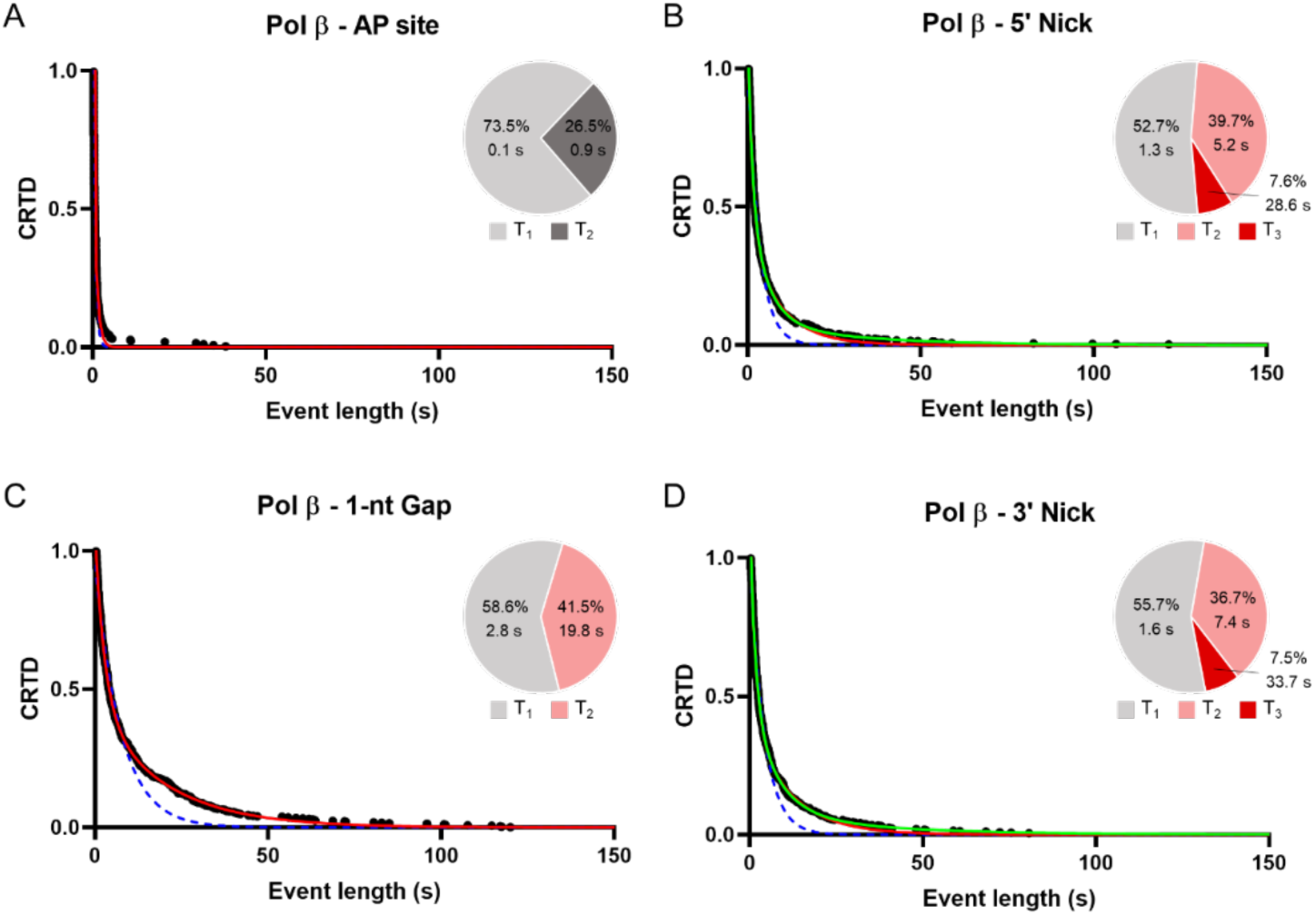
Pol β-DNA binary interactions. CRTD vs. time plot of binary complex lifetimes measured from Pol β binding events with DNA substrates containing an AP site **(A)**, 5’ nick **(B)**, 1-nt gap **(C)**, and 3’ nick **(D)**. Experimental data is represented by black circles. Single, double, and triple exponential decay fits are depicted as dashed blue, solid red, and solid green lines respectively. Pie charts show the relative fraction of molecules associated with the T_1_, T_2_, and if applicable T_3_ populations. The kinetic parameters and 95% confidence intervals described by the fits are shown in Supplemental Table 2.

### Characterization of the dynamic assembly of APE1-Pol β-5’ nick ternary complexes

According to the model of BER coordination between APE1 and Pol β, an APE1-Pol β-5’ nick ternary complex is formed before the dissociation of APE1 (12). To understand how these ternary complexes assemble, we used single molecule TIRF microscopy to record the order of APE1-Cy3 and Pol β-Cy5 binding to immobilized 5’ nick BER intermediates. A representative fluorescence trajectory of an APE1-Pol β-5’ nick ternary complex formation is shown in Figure 2. Of the 377 ternary complex formations that were observed, 263 (69.8%) formed when a preformed APE1-5’ nick binary complex was bound by Pol β (A^ON^-P^ON^) (Figure 5A). These events reflect the proper directionality of the BER pathway. In addition, we observed 102 ternary complexes (27.0%) which formed upon APE1 association to a preformed Pol β-5’ nick binary complex (P^ON^-A^ON^) and 12 ternary complexes (3.2%) which formed upon both APE1 and Pol β associating within the same frame (A/P^ON^) (Figure 5B-C). The low frequency of observed A/P^ON^ events relative to A^ON^-P^ON^ and P^ON^-A^ON^ events suggests that APE1-Pol β-5’ nick DNA ternary complexes primarily form through sequential binding of APE1 and Pol β. Thus, these results provide further evidence that APE1-Pol β-5’ nick ternary complexes primarily assemble through the independent association of each enzyme to the DNA substrate as opposed to the association of preformed APE1-Pol β complex.

**Figure 5.**
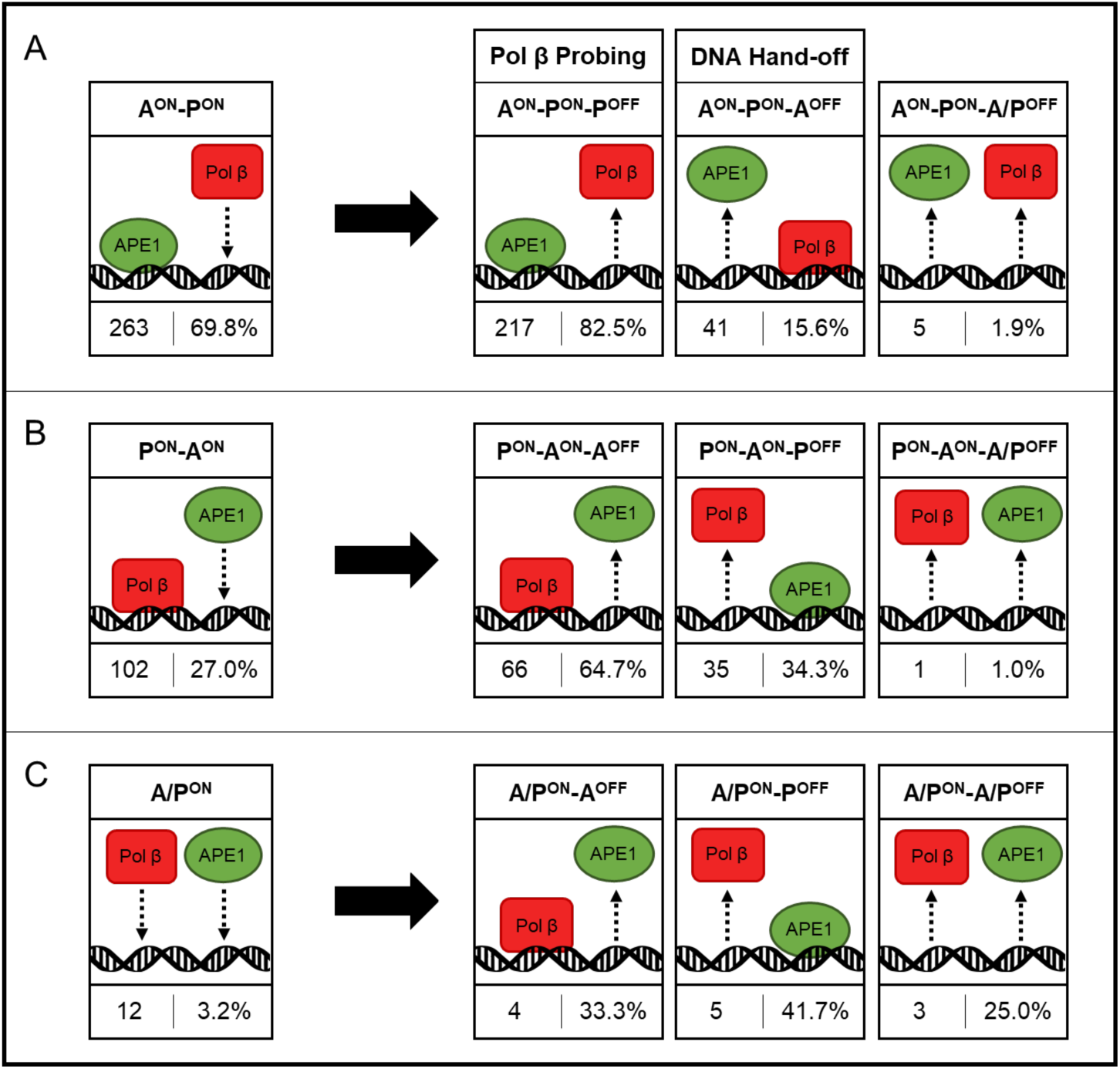
Ternary Complex Event Tree. **(A)** 263 ternary complexes formed when a preformed APE1-5’ nick binary complex was bound by Pol β (A^ON^- P^ON^), **(B)** 102 formed when a preformed Pol β-5’ nick binary complex was bound by APE1 (P^ON^-A^ON^), and **(C)** 12 formed when both proteins associated within the same 0.1 sec frame (A/P^ON^). The relative occurrence of each possible disassembly sequence (A^OFF^, P^OFF^, or A/P^OFF^) is displayed to the right of the black arrows. DNA hand-off events (A^ON^-P^ON^-A^OFF^) and Pol β probing events (A^ON^-P^ON^- P^OFF^) are specifically indicated.

The canonical BER coordination model proposes that after incision of the AP-site by APE1, the 5’ nick product is transferred to Pol β through the formation of an APE1-Pol β-5’ nick ternary complex. To determine the stability of the APE1-Pol β-5’ nick ternary complex, we measured the duration that each ternary complex remained intact (i.e., the ternary complex lifetime) for ternary events in which Pol β associated to a preformed APE1-5’ nick complex (A^ON^-P^ON^) (Figure 6A, inset). The average lifetime of these 263 ternary complexes was then quantified using CRTD analysis (Figure 6A). Fitting of the CRTD plot was best described by two kinetic processes, suggesting the existence of a short-lived (T_1_) and long-lived (T_2_) subpopulation of ternary complexes (Supplemental Table 3). The relative fraction of ternary complexes which dissociated according to the T_1_ and T_2_ processes is shown in Figure 6B. In the majority of ternary events (68.8%), the mean lifetime of the ternary complex was relatively short-lived (0.6 sec), while the remaining 31.2% of ternary events exhibited a longer-lived ternary complex (7.3 sec) (Supplemental Table 3). This finding led us to investigate if the short-lived and long-lived subpopulations may reflect different sequences of ternary complex disassembly.

**Figure 6.**
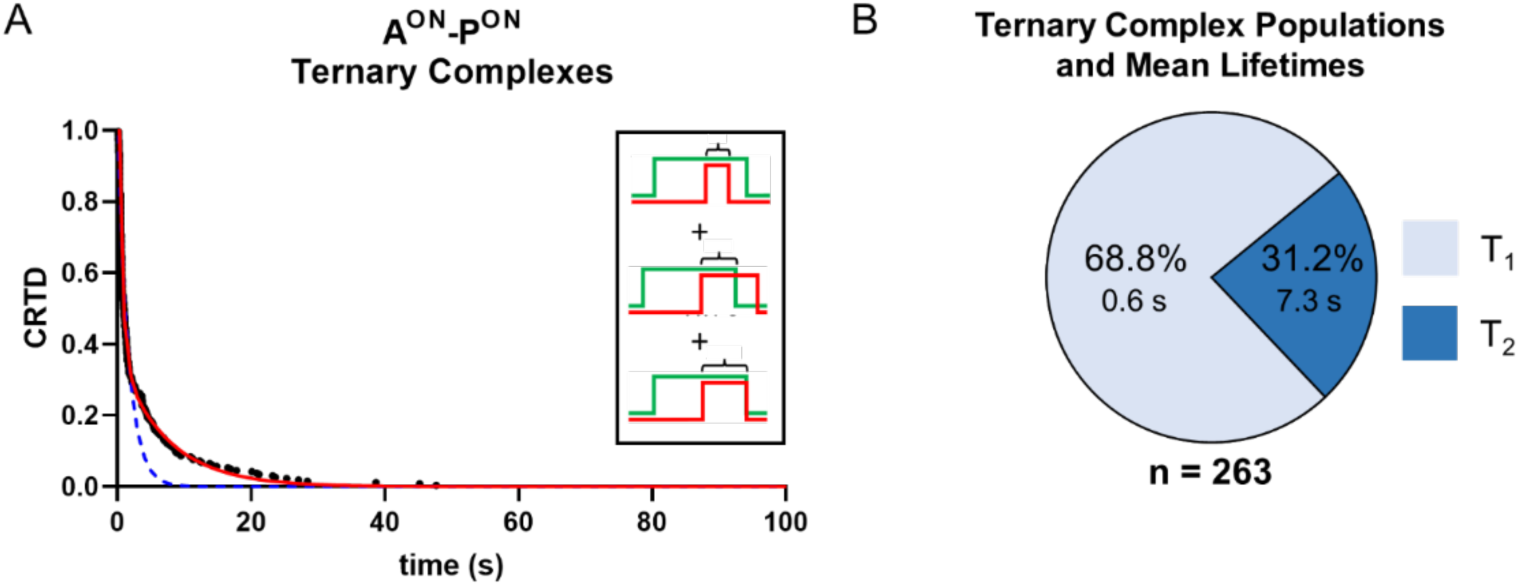
Analysis of A^ON^-P^ON^ ternary complex lifetimes. **(A)** CRTD vs. time for the lifetime of APE1-Pol β-5’ nick ternary complexes which formed when Pol β associated with a preformed APE1-5’ nick binary complex (A^ON^-P^ON^) (n=263). Black circles represent experimental data and single and double exponential fits are shown in dashed blue and solid red lines, respectively. (B) The fraction of ternary complexes which participate in the fast dissociating (T_1_) and slow dissociating (T_2_) processes is represented with light blue and dark blue segments of a pie chart, respectively. The kinetic parameters and 95% confidence intervals described by the fits are shown in Supplemental Table 3.

In order to distinguish whether the short-lived and long-lived populations in Figure 6 reflect different sequences of APE1 and Pol β dissociation, we first categorized ternary events based on the order of enzyme disassociation. Of the 263 A^ON^-P^ON^ ternary events, 217 events (82.5%) ended with a Pol β disassociation step (A^ON^-P^ON^-P^OFF^) (Figure 5A). These events, which we refer to as Pol β probing events, reflect unsuccessful transference of the DNA substrate. In many cases, after the formation of the APE1-5’ nick DNA complex, multiple repeated Pol β probing events were observed before the eventual dissociation of APE1. In contrast, 41 events (15.6%) ended with an APE1 disassociation step (A^ON^-P^ON^-A^OFF^). These events, which we refer to as DNA hand-off events, are true to the directionality of the BER pathway and reflect the transfer of the 5’ nick APE1 product from APE1 to Pol β. Only 5 ternary complexes (1.9%) ended with both APE1 and Pol β dissociating within the same 0.1 second frame window (A^ON^-P^ON^-A/P^OFF^). These rarely observed events correspond to either sequential dissociation events that are unresolved by the frame rate of the camera or represent the simultaneous dissociation of APE1 and Pol β from the immobilized 5’ nick DNA. These results provide evidence that APE1-Pol β-5’ nick ternary complexes primarily disassemble through the independent dissociation of each enzyme. Furthermore, these results suggest that the transfer of APE1’s product from APE1 to Pol β (DNA hand-off) can indeed occur under the conditions tested. Importantly, this is consistent with ensemble studies that showed APE1 to Pol β hand-off occurring on a similar time scale (11). However, association of Pol β to the APE1-5’ nick complex most frequently results in Pol β dissociation and unsuccessful transfer of the DNA (Pol β probing).

In addition to the 263 ternary complexes that formed upon Pol β association to an APE1-DNA complex (A^ON^-P^ON^), we observed 102 ternary complexes (27.0%) which formed upon APE1 association to a Pol β-5’ nick complex (P^ON^-A^ON^) (Figure 4.4B). Of these, 35 events (34.3%) ended with a Pol β disassociation step (P^ON^-A^ON^-P^OFF^). These events reflect the transfer of the 5’ nick APE1 product from Pol β to APE1. Another 66 events (64.7%) ended with an APE1 disassociation step (P^ON^-A^ON^-A^OFF^), which correspond to the association and subsequent dissociation of APE1 from Pol β-5’ nick complexes. The remaining 1 event (1.0%) ended with both proteins dissociating within the same frame (P^ON^-A^ON^-A/P^OFF^). These results provide further evidence that APE1-Pol β-5’ nick ternary complexes primarily assemble and disassemble through sequential association and dissociation of each enzyme. Furthermore, the finding that A^ON^-P^ON^ events primarily end with Pol β dissociation (A^ON^-P^ON^-P^OFF^), and P^ON^-A^ON^ events primarily end with APE1 dissociation (P^ON^-A^ON^-A^OFF^), indicates that the order of disassembly is dependent on the order in which the proteins assemble on the DNA substrate. This suggests that the initially bound protein forms more stable DNA interactions than the incoming protein. It’s likely that this reflects the site-specific DNA binding of the initially bound protein to the 5’ nick, which may obstruct accessibility for the incoming protein.

To better understand what may cause an APE1-Pol β-5’ nick ternary complex to result in DNA hand-off vs. Pol β probing, we measured the ternary complex lifetime of each DNA hand-off and Pol β probing event (Figure 7A). The average lifetime of ternary complexes for Pol β probing events (A^ON^-P^ON^-P^OFF^) and DNA hand-off events (A^ON^-P^ON^-A^OFF^) was then quantified using CRTD analysis (Figure 7B-C). Both the Pol β probing and DNA hand-off CRTD plots were best described by double exponential models reflecting two kinetic processes, suggesting the existence of a short-lived (T_1_) and long-lived (T_2_) population of ternary complexes in both cases (Supplemental Table 4). In the majority of Pol β probing events (71.0%), the mean lifetime of the ternary complex was relatively short-lived (0.4 sec), while the majority of DNA hand-off events (59.9%) exhibited a relatively longer-lived ternary complex (11.0 sec) (Figure 7D, Supplemental Table 4). These results indicate that after association of Pol β to the APE1-5’ nick complex, the ternary complex is more likely to end with DNA hand-off (APE1 dissociation) if Pol β remains bound to the APE1-5’ nick complex for an extended amount of time (∼11.0 sec). This suggests that the order of ternary complex disassembly (Pol β probing vs. DNA hand-off) is dependent on the disassociation of APE1 and the duration that Pol β remains bound near the APE1-5’ nick complex.

**Figure 7.**
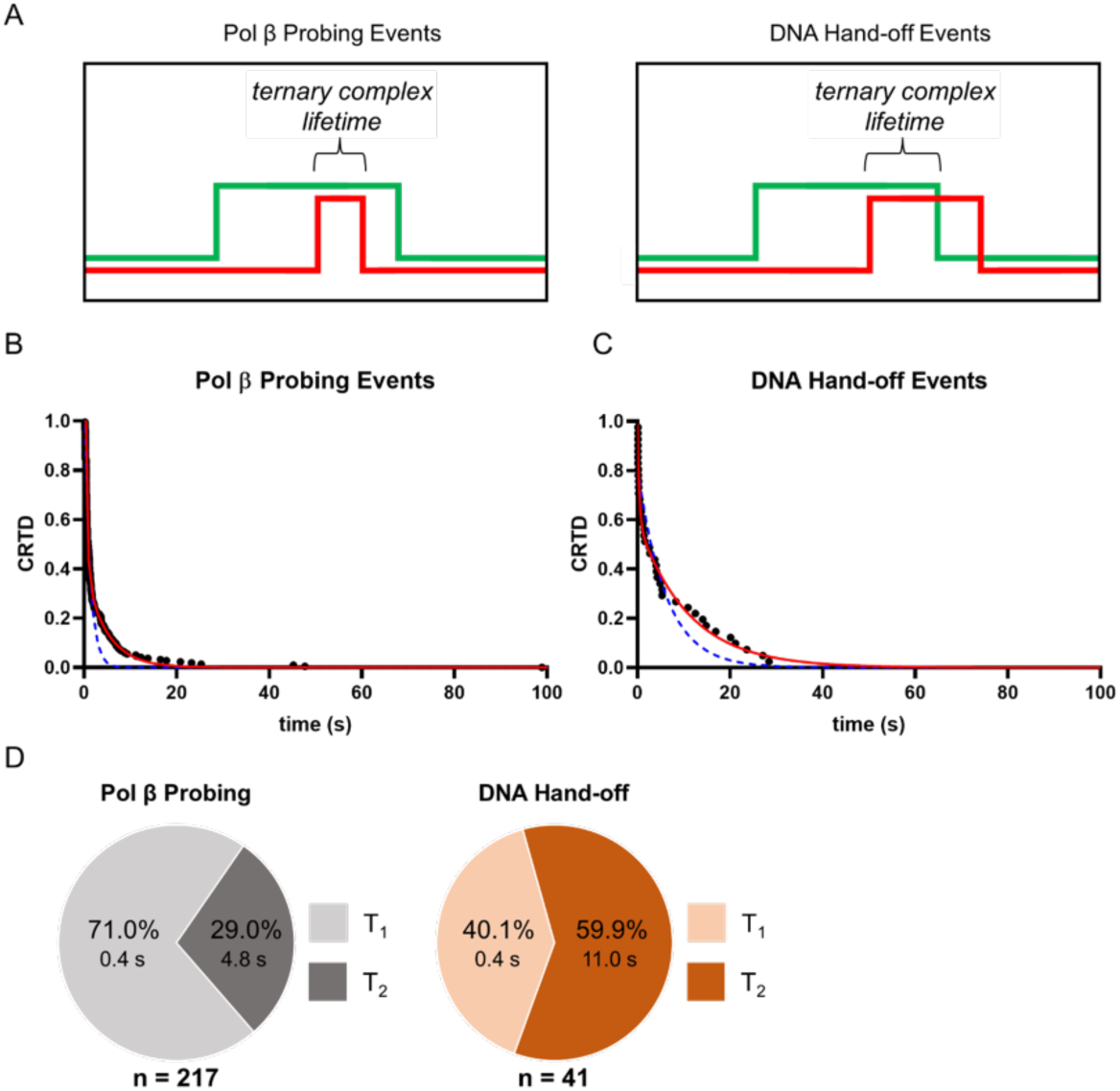
Lifetime of ternary complex in Pol β probing vs. DNA hand-off events. **(A)** The lifetime of a ternary complex is the duration that both proteins are simultaneously bound to the immobilized DNA. CRTD vs. time plots of ternary complex lifetimes measured from Pol β probing events **(B)** and DNA hand-off events **(C)**. Experimental data is represented by black circles and single and double exponential fits are depicted as dashed blue and solid red lines respectively. (D) The fraction of ternary complexes which participate in the fast dissociating (T_1_) and slow dissociating (T_2_) Poisson processes (Supplemental Table 4) observed for Pol β probing events (gray) and DNA hand-off events (orange), is represented with pie charts.

## DISCUSSION

In order to progress through the steps of BER, the correct BER enzyme should be bound to the BER intermediate at the relevant stage in the pathway. We found that APE1 and Pol β both form the most stable complexes with their respective DNA substrates and products compared to other BER intermediates in the pathway. Binary interactions between APE1 and the AP site BER intermediate had the longest mean lifetime compared to downstream BER intermediates (Figure 3, Supplemental Table 1). Similarly, Pol β exhibited the longest mean lifetime with its substrates and products: 5’ nick, 1-nt gap, and 3’ nick (Figure 4, Supplemental Table 2). Compared to APE1, Pol β formed much weaker complexes with BER intermediates that were not a canonical substrate or product of the enzyme (AP site and undamaged DNA). This is consistent with previous findings that indicate Pol β weakly associates with DNA substrates lacking a free 5’ phosphate (16). These findings are consistent with previous work which suggested that proper association of BER enzymes at the correct stage of BER could be explained, in part, by the target binding specificity inherent to each BER enzyme (16). Furthermore, lower affinity between BER enzymes and upstream or downstream BER intermediates in the pathway could minimize the competitive inhibition caused by the association of BER enzymes that are not relevant to the current stage of BER. Thus, the binding specificity of each BER enzyme for specific BER intermediates likely contributes to the proper directionality of BER, helping to ensure the correct proteins are bound at the proper BER step for efficient pathway progression.

Coordinated handling of the damaged DNA by BER enzymes between the steps of BER can minimize exposure of damaged DNA and the potential for further genomic instability (12,29). Previous studies have suggested a possible model of APE1-Pol β coordination where Pol β binds the flanking DNA of the APE1-5’ nick complex, promotes the displacement of APE1 and engages the 5’ nick (15,16,30). To explore whether Pol β displaces APE1 from the DNA, we observed the real-time formation of APE1-Pol β-5’ nick ternary complexes using single molecule TIRF microscopy. Our results indicate that Pol β dissociated from the APE1-5’ nick complex shortly after binding 82.5% of the time (i.e., Pol β probing events) (Figure 5A). The remaining ternary complexes that ended with APE1 dissociation resulted in DNA hand-off and on average lasted over 20 times longer (Figure 7D). These results indicate that the likelihood a given ternary event will end with DNA hand-off is largely dependent on how long Pol β remains bound near the APE1-5’ nick complex. This finding, along with the high frequency of observed Pol β probing events, suggests that Pol β does not efficiently evict APE1 from the 5’ nick under the conditions tested.

To efficiently locate target sites in the genome, Pol β utilizes a form of processive searching called DNA hopping (31,32). In the DNA hopping model, the DNA-binding protein dissociates to a point where all intermolecular bonds are broken, but since the protein is still very close to where it was originally bound it can re-associate with high probability to the same or nearby site on the DNA (33). Previous studies have indicated that Pol β uses this mechanism to “hop” along the length of DNA and can scan a region of ∼24 bases prior to fully disassociating from the DNA (31,32). In the context of the cell where there are long spans of DNA flanking the APE1-5’ nick complex, DNA hopping could allow Pol β to remain in the vicinity of the lesion, extending the amount of time that Pol β can wait for APE1 to dissociate from the 5’ nick. Thus, it is possible that frequent probing of the APE1-5’ nick complex by Pol β could be a means to capture the lesion immediately after APE1 dissociation.

Finally, while this model describes how APE1-Pol β coordination may occur independent of other BER factors, it’s possible that other factors might contribute to coordination efficiency that were not included here, such as the scaffolding protein X-ray repair cross-complementing 1 (XRCC1). XRCC1 harbors binding domains to multiple other DNA repair enzymes such as Lig3α (34), Pol β (35), APE1 (14), poly (ADP-ribose) polymerase 1 (PARP1) (36), and multiple DNA glycosylases (37,38). Many of these binding domains are spatially distinct, such as the binding domains for Pol β, Lig3α and APE1. This could suggest that XRCC1 may facilitate the formation of multimeric BER complexes, such as the transient complexes involved in BER coordination (39-41). It’s possible that XRCC1 could bridge APE1 and Pol β, keeping Pol β near the lesion long enough for APE1 to dissociate. Moreover, XRCC1 interactions with APE1 have been shown to promote APE1 product release (14). Thus, XRCC1 may act to promote APE1 release when Pol β is ready to accept the BER intermediate. The single molecule TIRF approach developed here could be extended to examine the effects of XRCC1 and other BER factors on the formation of ternary complexes with BER intermediates.

## SUPPLEMENTARY DATA

Supplementary Data are available online.

