## Supplemental Data for "Visualizing the Coordination of APE1 and DNA Polymerase β During Base Excision Repair"

| | | Analysis of APE1-DNA interactions | | | | | |
|---|---|---|---|---|---|---|---|
| | | $T_1$ Population | | $T_2$ Population | | $T_3$ Population | |
| BER intermediate | *N* | Pop (%) | $\tau_1$ (s) | Pop (%) | $\tau_2$ (s) | Pop (%) | $\tau_3$ (s) |
| Abasic DNA | 223 | 50.8 | 3.0 | 49.3 | 41.1 | | |
| *95% CI* | | *(49.9 – 51.7)* | *(2.9 – 3.1)* | | *(39.9 – 42.4)* | | |
| 5' Nick DNA | 570 | 38.8 | 1.2 | 46.2 | 8.3 | 15.1 | 36.5 |
| *95% CI* | | *(37.8 – 39.8)* | *(1.1 – 1.2)* | | *(7.7 – 8.8)* | *(12.6 – 17.8)* | *(32.1 – 42.3)* |
| 1 nt Gap DNA | 1151 | 21.8 | 1.2 | 48.3 | 4.3 | 29.9 | 30.9 |
| *95% CI* | | *(19.6 – 24.4)* | *(1.1 – 1.3)* | | *(4.1 – 4.6)* | *(29.0 – 30.7)* | *(30.0 – 31.9)* |
| 3' Nick DNA | 156 | 45.8 | 1.2 | 54.2 | 12.4 | | |
| *95% CI* | | *(44.6 – 47.1)* | *(1.2 – 1.3)* | | *(12.0 – 12.8)* | | |
| Undamaged DNA | 362 | 63.7 | 2.6 | 36.3 | 22.8 | | |
| *95% CI* | | *(62.7 – 64.7)* | *(2.6 – 2.7)* | | *(21.9 – 23.7)* | | |

**Supplemental Table 1** Binary complex mean lifetime for APE1 on BER intermediates.

N represents the total number of observed counts. $T_1$, $T_2$, and $T_3$ distinguish the independent Poisson processes. Pop is the coefficient of the exponential terms obtained from the fits and represents the relative fraction of all observed objects represented by the population. $\tau_i$ represent the mean lifetime of the respective population. The lower and upper bounds of the 95% confidence intervals (CI) are shown for each reported parameter.

| | | Analysis of Pol β-DNA interactions | | | | | |
|---|---|---|---|---|---|---|---|
| | | $T_1$ Population | | $T_2$ Population | | $T_3$ Population | |
| BER intermediate | *N* | Pop (%) | $\tau_1$ (s) | Pop (%) | $\tau_2$ (s) | Pop (%) | $\tau_3$ (s) |
| Abasic DNA | 258 | 73.5 | 0.1 | 26.5 | 0.9 | | |
| *95% CI* | | *(67.8 - 84.3)* | *(0.1 – 0.2)* | | *(0.8 – 1.2)* | | |
| 5' Nick DNA | 833 | 52.7 | 1.3 | 39.7 | 5.2 | 7.6 | 28.6 |
| *95% CI* | | *(49.8 – 55.3)* | *(1.2 – 1.3)* | | *(4.7 – 5.8)* | *(5.4 – 10.4)* | *(21.6 – 41.0)* |
| 1 nt Gap DNA | 639 | 58.6 | 2.8 | 41.5 | 19.8 | | |
| *95% CI* | | *(57.9 – 59.2)* | *(2.8 – 2.8)* | | *(19.4 – 20.1)* | | |
| 3' Nick DNA | 546 | 55.7 | 1.6 | 36.7 | 7.4 | 7.5 | 33.7 |
| *95% CI* | | *(52.4 – 58.5)* | *(1.5 – 1.7)* | | *(6.1 – 8.6)* | *(3.9 – 13.1)* | *(22.5 – 65.3)* |

**Supplemental Table 2** Binary complex mean lifetime for Pol β on BER intermediates

N represents the total number of observed counts. $T_1$, $T_2$, and $T_3$ distinguish the independent Poisson processes. Pop is the coefficient of the exponential terms obtained from the fits and represents the relative fraction of all observed objects represented by the population. $\tau_i$ represent the mean lifetime of the respective population. The lower and upper bounds of the 95% confidence intervals (CI) are shown for each reported parameter.

| | | Analysis of $A^{ON}$-$P^{ON}$ ternary complex lifetimes | | | |
|---|---|---|---|---|---|
| | | $T_1$ Population | | $T_2$ Population | |
| Event Type | *N* | Pop (%) | $\tau_1$ (s) | Pop (%) | $\tau_2$ (s) |
| $A^{ON}$-$P^{ON}$ Ternary Complex | 263 | 68.8 | 0.6 | 31.2 | 7.3 |
| *95% CI* | | *(66.7 – 70.8)* | *(0.6 – 0.6)* | - | *(6.5 – 8.2)* |

**Supplemental Table 3 Ternary complex mean lifetime for APE1-Pol β-5' nick ternary complex formations.**

*N* represents the total number of observed counts. $T_1$ and $T_2$ distinguish the two identified independent Poisson processes. Pop is the coefficient of the exponential terms obtained from the fits and represents the relative fraction of all observed objects represented by the subpopulation. $\tau_i$ represent the mean lifetime of the respective population. The lower and upper bounds of the 95% confidence intervals (CI) are shown for each reported parameter.

| | | Analysis of ternary complex event lifetimes | | | |
|---|---|---|---|---|---|
| | | $T_1$ Population | | $T_2$ Population | |
| Event Type | *N* | Pop (%) | $\tau_1$ (s) | Pop (%) | $\tau_2$ (s) |
| Pol β probing | 217 | 71.0 | 0.4 | 29.0 | 4.8 |
| *95% CI* | | *(68.9 – 73.0)* | *(0.4 – 0.5)* | - | *(4.3 – 5.4)* |
| DNA hand-off | 41 | 40.1 | 0.4 | 59.9 | 11.0 |
| *95% CI* | | *(35.6 – 44.5)* | *(0.2 – 0.7)* | - | *(9.4 – 13.1)* |

**Supplemental Table 4** Ternary complex mean lifetime for Pol β probing and DNA hand-off events.

*N* represents the total number of observed counts. $T_1$ and $T_2$ distinguish the two identified independent Poisson processes. Pop is the coefficient of the exponential terms obtained from the fits and represents the relative fraction of all observed objects represented by the subpopulation. $k_{d,i}$ represents the dissociation rate constant and $\tau_i$ represent the mean lifetime of the respective population. The lower and upper bounds of the 95% confidence intervals (CI) are shown for each reported parameter.
